# FGF21 regulates melanogenesis in alpaca melanocytes *via* ERK1/2-Mediated MITF downregulation

**DOI:** 10.1101/147082

**Authors:** Ruiwei Wang, Tianzhi Chen, Bingling Zhao, Ruiwen Fan, Kaiyuan Ji, Xiuju Yu, Xianjun Wang, Changsheng Dong

## Abstract

Fibroblast growth factor 21 (FGF21) is known as a metabolic regulator to regulate the metabolism of glucose and lipids. However, the underlying mechanism of FGF21 on melanin synthesis remains unknown. Therefore, the current study investigates the effect of FGF21 on melanogenesis in alpaca melanocytes. We transfected the FGF21 into alpaca melanocytes, then detected the melanin contents, protein and mRNA levels of pigmentation-related genes in order to determine the melanogenesis-regulating pathway of FGF21. The results showed that FGF21 overexpression suppressed melanogenesis and decreased the expression of the major target genes termed *microphthalmia-associated transcription factor* (MITF) and its downstream genes, including tyrosinase (TYR) and tyrosinase-related protein 2 (TRP2). However FGF21 increased the expression of phospho-extracellular signal-regulated kinase (p-Erk1/2). In contrast, FGF21-siRNA, a small interference RNA mediating FGF21 silencing, abolished the inhibition of melanogenesis. Altogether, FGF21 may decrease melanogenesis in alpaca melanocytes via ERK activation and subsequent MITF downregulation, which is then followed by the suppression of melanogenic enzymes and melanin production.

## 1. Introduction

Melanin, synthesized by melanocytes, determines the color of fur, hair and skin. Melanin containing melanocyte-specific organelles, known as melanosomes, were transfered from melanocytes to adjacent keratinocytes, which formed the distribution of pigmentation and eventually variability of visible colors^[1]^. The melanins are produced in melanocytes and have two main types: brown/black, eumelanin, and red/yellow, pheomelanin^[2]^. It has been confirmed that the melanin synthesis pathway is closely related to melanocyte-specific enzymes - tyrosinase, tyrosinase-related protein 1, and tyrosinase-related protein 2^[3]^. Among these enzymes, tyrosinase is the first rate-limiting enzyme^[4]^. The upregulation of tyrosinase contributes to the formation of eumelanin^[5]^, whereas its downregulation produces pheomelanin^[6]^. The transcriptional regulation of the three major melanogenic enzymes are regulated by MITF^[7]^, which acts as a central role for melanocyte development, survival, proliferation and melanogenesis^[8]^.

FGF21 showed an abnormal expression in murine melanoma cells^[9]^. Thus the current study looks at whether or not FGF21 is involved in melanogenesis. MITF expression was inhibited by activating FGF/FGFR/MEK/ERK signaling pathways^[10]^. FGF21 is a new number of the FGF super family^[11]^. This study investigated if FGF21 would also similar to FGFR/MEK/ERK signaling pathways, which would lead to MITF downregulation. A previous study showed that FGF receptor mechanisms mediate FGF21 signal pathways^[12]^, which results in FGF21-FGFR complex induced ERK cascade and sustained activation of ERK^[13]^. The ERK signaling pathway is involved in melanin synthesis^[14]^, activation of ERK phosphorylates MITF at serine 73^[15]^, followed by degradation of MITF^[16]^. Therefore, FGF21 might control melanogenesis of alpaca melanocytes via ERK1/2 signal pathway. In this study, FGF21 was selected as a candidate gene for melanogenesis regulation.

## 2. Results

### 2.1. Effects of FGF21 on melanogenesis in alpaca melanocytes

To detect the effect of FGF21 on melanogenesis in aplaca melanocytes, alpaca melanocytes were transfected with pMSCV-GFP vector encoding FGF21 and FGF21-siRNA. There are two experiments. The first looks at pMSCV-GFP-FGF21 and the pMSCV-GFP control group. The second looks at FGF21-siRNA and the corresponding control group. The qRT-PCR and western blot results showed that the relative mRNA and protein levels of FGF21 in the PMSCV-FGF21 group were 17.58 and 1.82 times higher than that in the control group, respectively (Figure 1A-C). However, in the FGF21-siRNA transfected group, the expression of FGF21 reduced by 4.10 times and 0.62 times for mRNA and protein levels compared with its control group, respectively (Figure 2D-F). These results showed that pMSCV-FGF21 and FGF21-pMSCV has been transfected into alpaca melanocytes efficiently.

**Figure 1.**
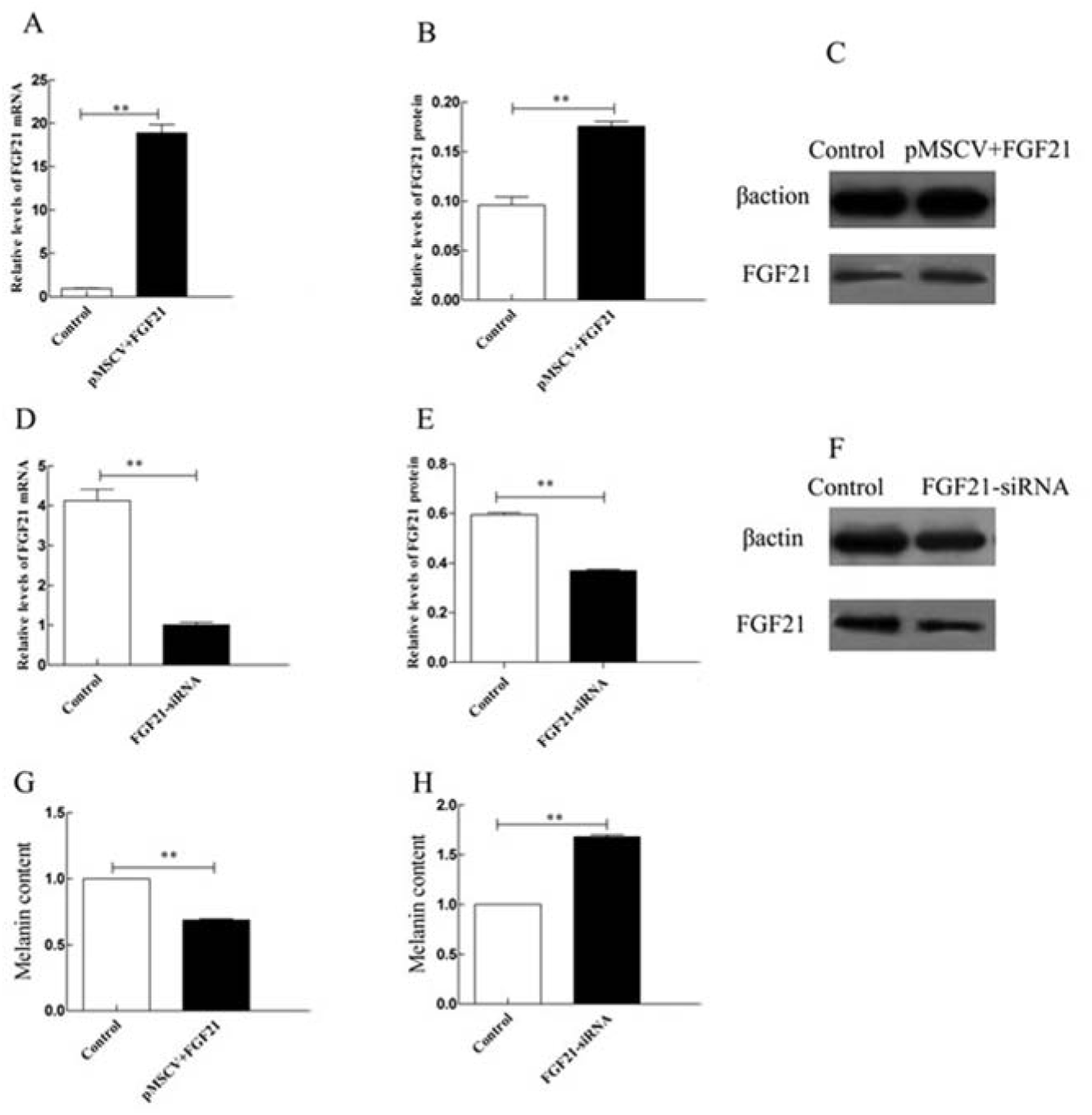
Effect of FGF21 on melanogenesis in alpaca melanocytes. Cells were transfected with pMSCV-FGF21 and FGF21-siRNA as well as their corresponding controls. (A-F) The expression levels of *FGF21* were analyzed by qRT-PCR and western blot. (G), (H) Melanin contents were measured, as described in the Materials and Methods section. Results are shown as the mean ± SD (*n* = 3 each), ** *p* < 0.01.

**Figure 2.**
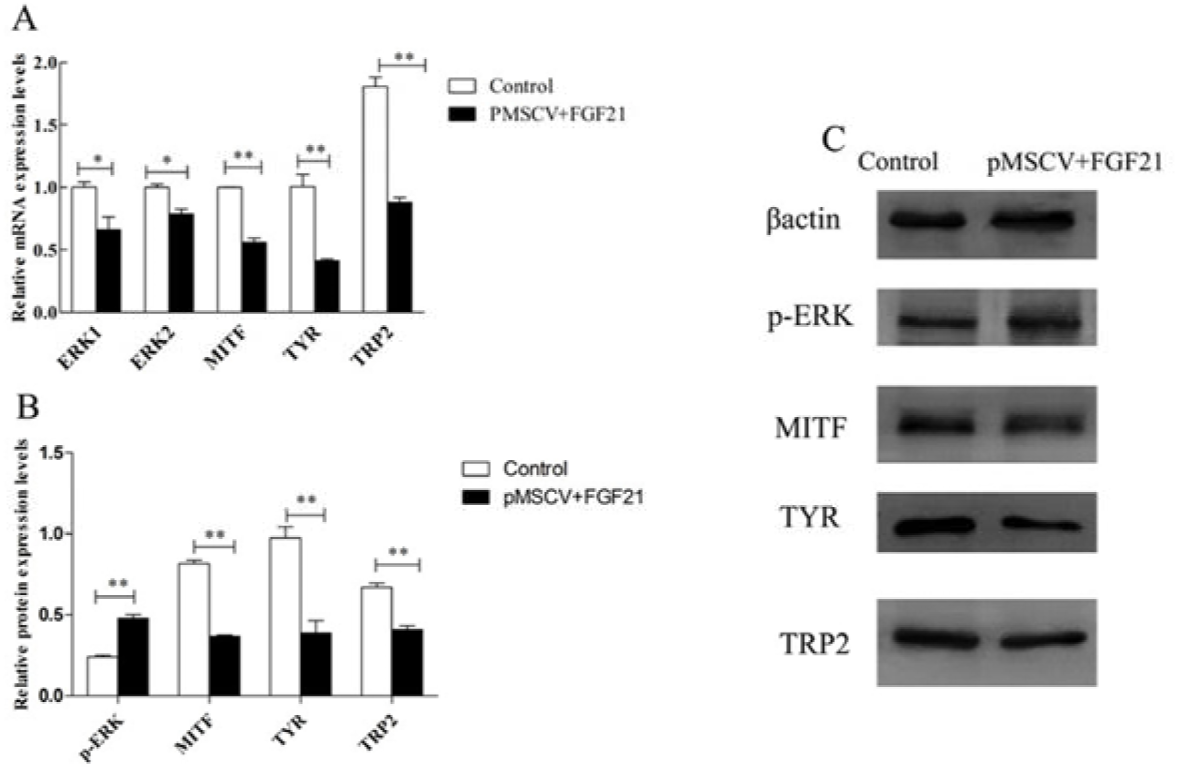
FGF21 overexpression affects melanin synthesis in alpaca melanocytes. The pMSCV-FGF21 and pMSCV-GFP was transfected into the alpaca melanocytes effectively. Total RNA and protein was extracted for the analysis of qRT-PCR and western blot. (A) The mRNA expression of ERK1/2, MITF, TYR, TRP2 was amplified with specific primers and measured by qRT-PCR, and 18S was used as an mRNA loading control. (B), (C) Western blot of cells for relative protein abundance with antibodies recognized phospho-specific ERK, MITF, TYR and TRP2. Protein loading was normalized relative to β-actin. The results are shown as the mean ± SD (*n* = 3 each), * *p* < 0.05, ** *p* < 0.01.

To confirm the effect of FGF21 on melanogenesis in aplaca melanocytes, the amount of melanin contents were measured. As shown in Figure 1G, the melanin contents decreased significantly in alpaca melanocytes transfected with pMSCV-FGF21-GFP compared to pMSCV-GFP group, while alpaca melanocytes transfected with FGF21-siRNA produced more melanin contents (Figure 1H).

### 2.2. FGF21 induces MITF downregulation and activates Erk1/2 signaling pathways

In alpaca melanocytes, overexpression of FGF21 reduced melanin content, thus FGF21 may have effects on MITF expression, which functions as a key regulator on melanogenesis^[17]^.As shown in Figure 2A-C, qRT-PCR and western blotting results showed that, for the FGF21-transfected group, the relative level of MITF were substantially lower for mRNA and protein when compared with the control group.

Previous studies have demonstrated that sustained phosphorylation of ERK degrades MITF and subsequent MITF phosphorylation^[18]^. Therefore, this study will be verified if FGF21 activates ERK signaling pathways. The qRT-PCR and western blot results showed that FGF21 stimulated the ERK1/2 signaling pathway. Moreover, the phosphorylation of MITF was associated with activation of ERK, leading to downregulation of TYR, TRP1 and TRP2^[19]^. In this study, the target genes of MITF and key melanic enzymes TYR and TRP2 for melanin production were also reduced by different amounts (Figure 2A-C). The results supported that FGF21 overexpression induced ERK signaling pathways and downregulation of MITF, TYR and TRP2, which resulted in a reduction of melanin contents.

### 2.3. FGF21-siRNA abolishes hypopigmentation via inhibiting ERK1/2 signaling pathway

The effect of FGF21-siRNA on ERK1/2 signaling and melanogenesis was further investigated. FGF21-siRNA abrogated the activation of ERK1/2 induced by FGF21 was found. A previous study showed that ERK1/2 inhibitor lead to the activation of melanin synthesis^[20]^. In this study, the relative expression of melanin contents in FGF21-siRNA-treated melanocytes increased significantly when compared with the control group. Collectively, the expression of MITF, TYR and TRP2 reversed at different levels after FGF21-siRNA transfection. The results showed that FGF21-siRNA may abolish hypopigmentation via inhibiting ERK1/2 signaling pathway.

**Figure 3.**
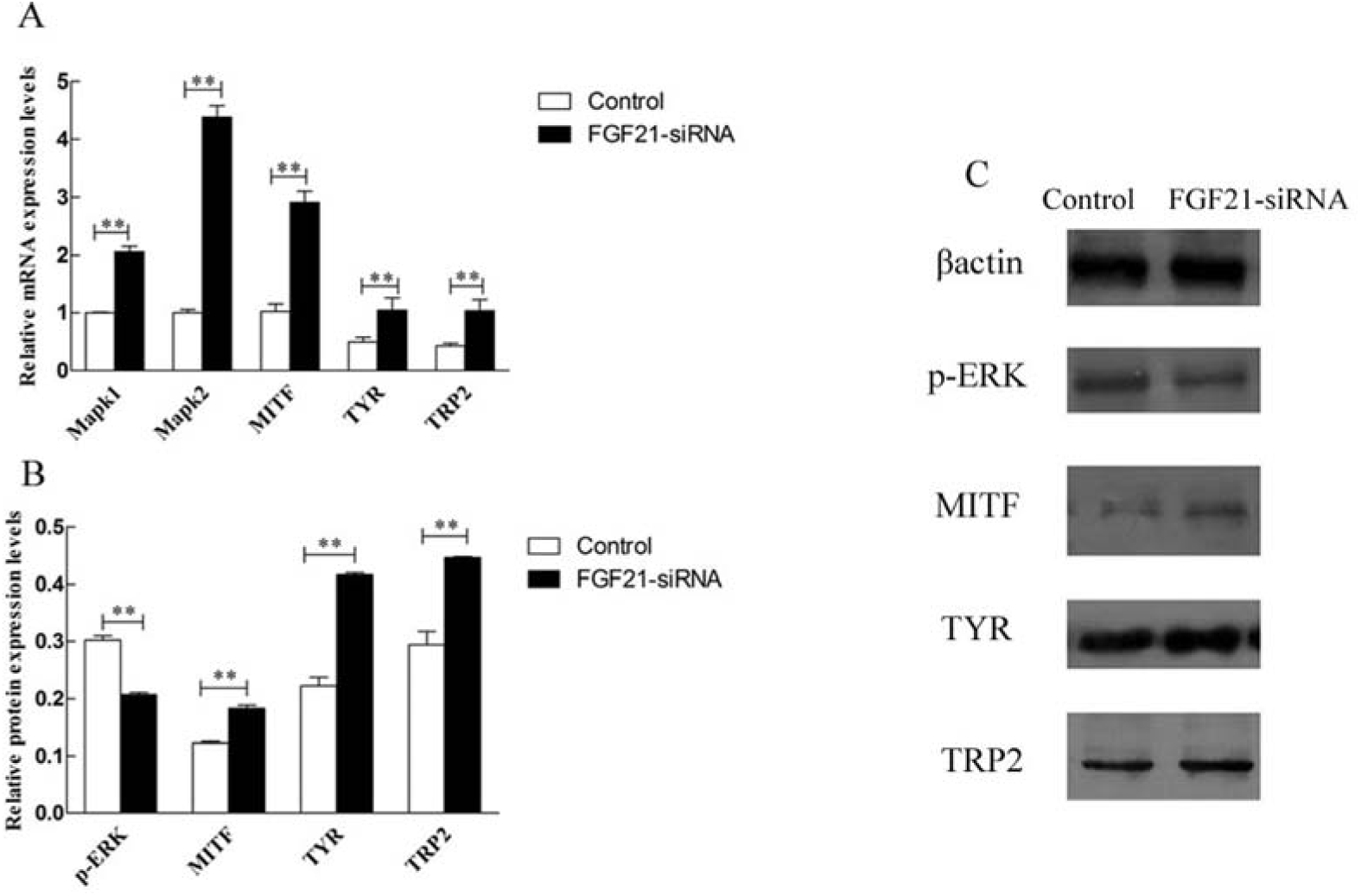
Effects of FGF21-siRNA on ERK1/2 pathway and melanogenesis-related genes. (A) The mRNA expression of FGF21 and melanogenesis-related genes were analyzed by qRT-PCR. (B) The protein expression of FGF21 and melanogenesis-related genes were analyzed by western blot. The results are shown as the mean ±SD (*n* = 3 each), \*\**p* < 0.01.

## 3. Discussion

Melanin is actually an indole polymer originated from the oxidation of tyrosine and tyrosinase-related enzymes^[21]^. TYR oxidizes L-tyrosine to 3,4-dihydroxy phenylalanine (DOPA) and then the oxidation of ODPA to dopaquinone^[22]^. TRP2, the dopachrome delta-isomerase (DCT), promotes DC to DHTCA^[23]^. With the catalysis of TRP1, eumelanin monomer named 5,6-dihydroxyindole-2-carboxylic acid (DHICA) composed eumelanin polymer^[24]^. MITF is a basic helix-loop-helix–leucine zipper (bHLH-LZ) transcription factor^[25,26]^, which regulates melanogenesis by binding to M-and E-box’s promoter of TYR, TRP1 and TRP2^[27,28]^. MITF is subjected to complex regulatory controls by multiple signal pathways, such as Wnt signaling pathway, cAMP pathway, extracellular signal-regulated kinase1/2 (ERK1/2) pathway, NO-mediated pathway regulated MITF at transcriptional or post-transcriptional levels^[29]^. ERK signaling cascade is one of the classic signal pathways, having an essential role in cell growth, development, proliferation and differentiation^[30]^. Increased activation of ERK pathway stimulated MITF for ubiquitin-dependent degradation^[31]^.

More and more biological functions of FGF21 were exhibited, but a comprehensive understanding of FGF21 on melanogenesis has been lacking. Several studies have shown that FGF21 could activate ERK1/2 pathways^[32]^. After alpaca melanocytes were transfected with PMSCV-FGF21, the protein, phospho-ERK1/2, increased and the mRNA, ERK1/2, decreased. The results show that FGF21 stimulates phosphorylation of Erk1/2 and activates Erk1/2 signal pathways. Previous studies have reported that phospho-Erk1/2 targets MITF phosphorylation and ubiquitin-dependent proteolysis^[33]^. The activation of ERK also had inhibitory effect on tyrosinase activity and transcription^[34,35]^. As expected, FGF21 decreased melanogenesis by a downregulation of MITF, TYR, TRP2, which might be associated with ERK activation.

However, the inhibition of ERK stimulated melanin synthesis and tyrosinase activity^[36]^. Alpaca melanocytes treated with FGF21-siRNA and its corresponding control group reversed the inital inhibitory effect of FGF21 on MITF and tyrosinase enzymes,. The augmented melanin pigmentation associated with upregulation of MITF is mediated by the inhibition of the ERK1/2 pathway.

## 4. Materials and Methods

### 4.1. Antibodies

Antibodies recognizing phospho-specific ERK1/2 (Thr202/Tyr204) were purchased from Cell Signaling Technology (Danvers, MA, USA). FGF21, MITF, TYR, TYRP2 were obtained from Abcam Biotechnology (Abcam, Cambridge, UK). (β-action was obtained from CW Biotech.

### 4.2. Cell cultures And Cell transfection

All alpaca melanocytes used in this study were established in Laboratories of Alpaca Biology, College of Animal Science and Technology, Shanxi Agricultural University, China. Alpaca melanocytes were cultured in Melanocyte Medium (MelM) (ScienCell Research Laboratories, Carlsbad, CA, USA), and were supplemented with 1% Melanocyte growth supplement (MELGS), 0.5% fetal bovine serum(FBS), 100u/ml Penicillin and 100ug/ml Streptomycin at 37□ in a 5% CO2-containing atmosphere.

FGF21 was synthesized and cloned into pMSCV-GFP plasmid (Addgene, Cambridge, MA, USA), resulting in the introduction of FGF21. FGF21-siRNA was purchased from Sangon Biotech, and mediates gene FGF21 silencing. Alpaca melanocytes were transfected using Lipofectamine 3000 reagent (Invitrogen, USA), according to the operation manual kit. 48h later, the melanocytes were harvested.

### 4.3. Melanin Content Assay

Melanin was extracted for a melanin content assay. Melanocytes were harvested and cleaned with cold PBS 3 times. Then cell precipitation was dissolved in 1 mol/L NaOH at 80□ for 30 mins. The relative melanin content (ug/10^6^cells) was measured at 475 nm using a Microplate Reader. Before measuring, melanocytes were observed and counted.

### 4.4. Total RNA Extraction and qRT-PCR Analysis

Total RNA was isolated from melanocyte cells using TRIzol Reagent (Life Technology). Synthesis of cDNA for real-time PCR analysis of FGF21 expression in cultured melanocytes was performed using the miRNA qRT-PCR kit according to the manufacturer’s instructions. The levels of gene mRNA were measured using SYBR®Premix Ex TaqTMII (Tli RNaseH Plus) according to the manufacturer’s instructions (TAKARA, Dalian, China).

Conditions for real-time PCR were as follows: preheating at 95□ for 5min, followed by 40 cycles of shuttle heating at 95□ for 30s, annealing temperature at 60□ for 30s, extending temperature at 72□ for 20s. Quantification of mRNA level for the target gene was performed using the comparative threshold cycle (CT) method. The expression of mRNA for the target genes was normalized relative to the abundance of 18S rRNA. The primer sequences are shown in Table 1.

**Table 1.**
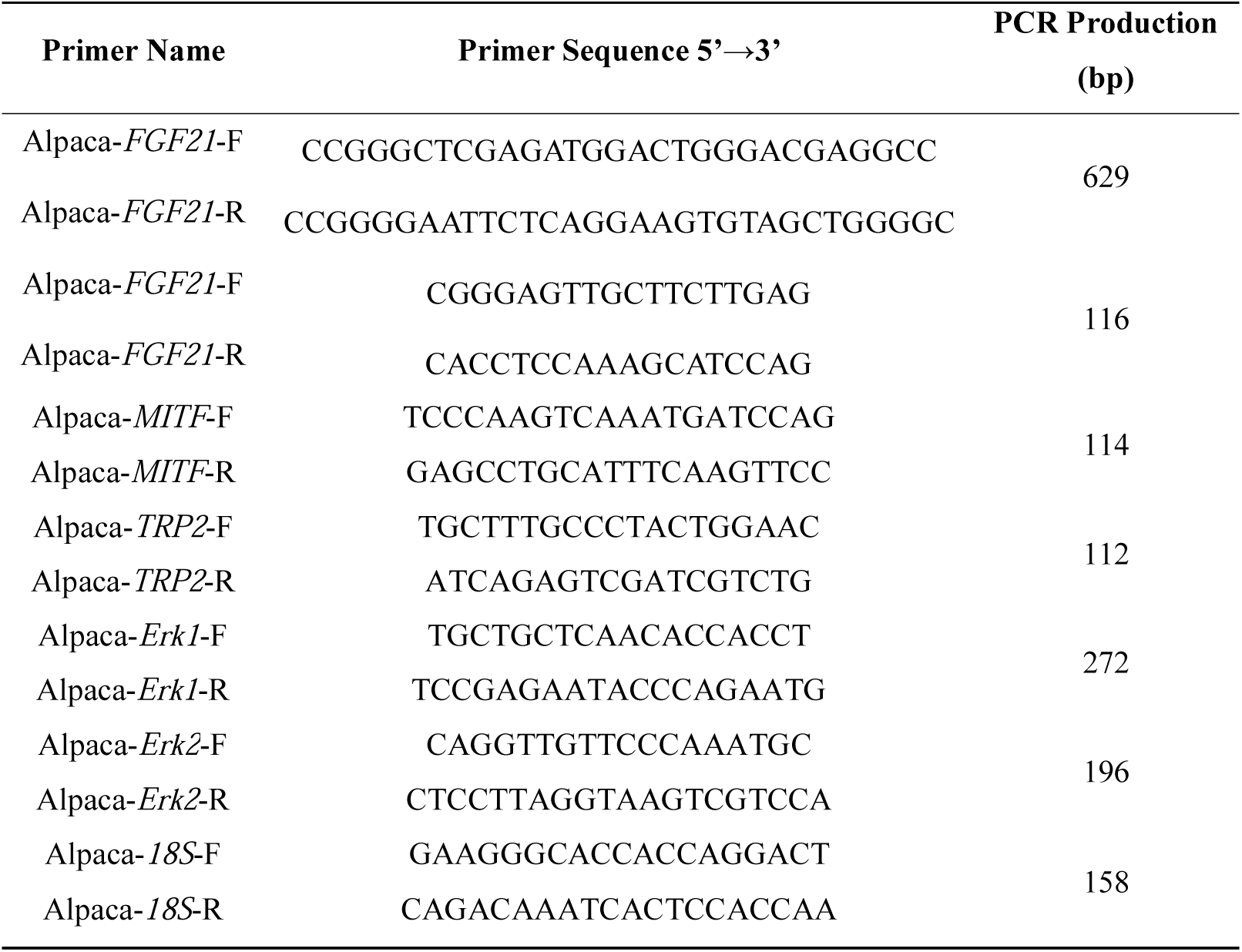
Primer sequences and qRT-PCR production for targeting gene

### 4.5. Western Blotting

Total protein was extracted from melanocytes using a protein extraction reagent (RIPA Lysis Buffer, Beyotime) according to the manufacturer’s guidelines. Concentration was measured using a nucleic acid/protein analyzer. Protein(200ng/lane) was separated using 10% SDS-PAGE electrophoresis and transferred onto nitrocellulose filter membranes. The membranes were blocked in 5% skim milk for 1 h. Then incubated membranes were diluted in 1×TBS with 0.1% Tween 20 at 4°C while being gentle shaken overnight. The next day, the membrane was washed three times with TBST and incubated with diluted HRP-conjugated secondary antibody for 1 h at 37°C. Finally, the membranes were washed 6 times with TBST for 5 minutes each. The protein abundance was analyzed using Image-Pro Plus Software, version 6.0 (Media Cybernetics), and normalized relative to β-actin in each lane.

### 4.6. Statistical analysis

All of the results were repeated in triplicate and expressed as mean ± standard errors. Each set of data was analyzed by GraphPad Prism 5.0 software (GraphPad Software Inc,CA, USA). P-values < 0.05 for the experimental and corresponding control groups were considered statistically significant, and p-values < 0.001 represent highly significant results.

## 5. Conclusions

In this research, we found FGF21 overexpression reduced melanin synthesis via ERK signaling pathway, accompanied with downregulation of MITF and target genes, including TYR and TRP2. In contrast, FGF21-siR stopped hypopigmentation by inhibiting ERK1/2, and the expression of MITF and melanin-specific enzymes increased when compared with the control groups. As a result, FGF21 may decrease melanogenesis of alpaca melanocytes via ERK1/2-Mediated MITF degradation. Altogether, FGF21 may decrease melanogenesis of alpaca melanocytes via ERK1/2-Mediated MITF downregulation.

## Acknowledgments

This research was supported by the High Technology Research and Development Program of China (863 Program) (2013AA102506), the Special Foundation for

Agro-scientific Research in the Public Interest (201303119) and the Aid Program for Innovative Research Team in Shanxi Agricultural University (CXTD201201).

